# Genome-wide CRISPR screen reveals genetic modifiers of Ca^2+^-mediated cell death

**DOI:** 10.1101/2023.01.13.523980

**Authors:** Oscar E. Reyes Gaido, Kate L. Schole, Mark E. Anderson, Elizabeth D. Luczak

## Abstract

Ca^2+^ is a fundamental determinant of survival in living cells. Excessive intracellular Ca^2+^ causes cellular toxicity and death but the genetic pathways contributing to Ca^2+^ induced cell death are incompletely understood. Here, we performed genome-wide CRISPR knock-out screening in human cells challenged with the Ca^2+^ ionophore ionomycin and identified genes and pathways essential for cell death after Ca^2+^ overload. We discovered 115 protective gene knockouts, 82 of which are non-essential genes and 21 of which belong to the druggable genome. Notably, members of store operated Ca^2+^ entry (SOCE), very long-chain fatty acid synthesis, and SWItch/Sucrose Non-Fermentable (SWI/SNF) pathways provided marked protection against Ca^2+^ toxicity. These results reveal pathways previously unknown to mediate Ca^2+^-induced cell death and provide a resource for the development of pharmacotherapies against the sequelae of Ca^2+^ overload in disease.

## INTRODUCTION

Ca^2+^ is a ubiquitous signaling ion in living cells and intracellular Ca^2+^ regulation has been coupled to various vital processes including proliferation, muscle contraction, fertilization, immune activation, neurotransmitter release, and vascular function *(1–8)*. Thus, cytosolic and organellar Ca^2+^ flux is tightly regulated. Ca^2+^ is typically maintained at a low concentration (10-100 nM) within cells, tens of thousands of times lower than the extracellular space (1-2 mM) *(2, 9)*. This intracellular concentration can transiently increase ∼10-fold as Ca^2+^ is used as a secondary signaling messenger *(2)*. Failure to maintain this homeostasis—for example, due to membrane instability or energetic failure—allows supraphysiologic concentrations to enter cells and trigger various pro-cell death pathways *(10)*.

Ca^2+^ is known to modulate several modes of cell death including apoptosis, necrosis, autophagic death, and anoikis *(10)*. Noteworthy known mechanisms of injury include *i)* mitochondrial permeability transition, where intracellular Ca^2+^ is internalized by mitochondria and triggers mitochondrial dysfunction and rupture *(11), ii)* calpains and caspases, pro-death proteases that require Ca^2+^ for induction *(12–14)*, and *iii)* Ca^2+^/Calmodulin-dependent protein kinase II (CaMKII) hyperactivity—which leads to inflammation and activation of apoptotic and necroptotic programs *(15–19)*. These pathways have become therapeutic targets with the hope to prevent tissue death. However, targeting of known Ca^2+^-dependent pathways to prevent tissue death has yet to be translated into clinical therapies due to several challenges. For example, myocardial ischemia/reperfusion injury is a major source of morbidity and well known to feature both Ca^2+^ overload and mitochondrial permeability transition *(20)*. While it is established that mitochondrial permeability transition induces cell death, the identity of the pore remains unclear and putative inhibitors of mitochondrial permeability (e.g. Cyclosporine A) have failed in clinical trials *(21, 22)*. Similarly, caspase, calpain, and CaMKII inhibitors have been long sought to prevent cell death in ischemic, septic, and traumatic injury but adverse effects, lack of selectivity, and lack of efficacy have precluded their use in patients *(19, 23–27)*. These examples highlight that our understanding of Ca^2+^ in cell death is incomplete and that identifying other targetable mediators of Ca^2+^ toxicity remains as a critical unmet need.

In the last decade, clustered regularly interspaced short palindromic repeats (CRISPR) and its associated nuclease Cas9 have enabled unprecedented tractability of genomic editing *(28, 29)*. Gene knockout can be achieved within days with high specificity. Pooled CRISPR viral libraries, which consist of a mixture of lentiviral particles where each virus encodes for a single gene knock-out, enable feasible interrogation of the entire genome in a single experiment *(30, 31)*. While this approach has been successful in identifying mediators of cell death in various contexts—including metabolic failure, hypoxia, infection, and senescence *(32–36)*—there remains a need for an unbiased examination of modifiers of cell survival in the context of excessive intracellular Ca^2+^.

In the present study, we modeled Ca^2+^ toxicity in an immortalized human cell line and performed genome-wide knockout perturbation using a CRISPR library. We identified 115 genes whose knockout allowed cells to survive Ca^2+^ overload, and 9 genes which sensitized cells to Ca^2+^. The genes outlined here will serve as a foundational resource for the elucidation of Ca^2+^-associated mechanistic effectors and for the development of therapeutic inhibitors.

## RESULTS

### CRISPR-screen identifies genetic modifiers of Ca^2+^ toxicity

To induce intracellular Ca^2+^ overload in human cells, we treated K562 cells with ionomycin—a Ca^2+^ ionophore. 24-hour ionomycin treatment was sufficient to induce cell death (Fig. 1A). This effect was dependent on the concentration of both ionomycin and Ca^2+^ present in the media; and physiological plasma Ca^2+^ concentration of 2.4 mM enabled near complete lethality at 10 µM ionomycin (Fig. 1A). To ensure toxicity is due to Ca^2+^ and not due to general ionomycin toxicity, we chelated Ca^2+^ in the media with EGTA which prevented ionomycin toxicity (Fig. 1B).

**Figure 1.**
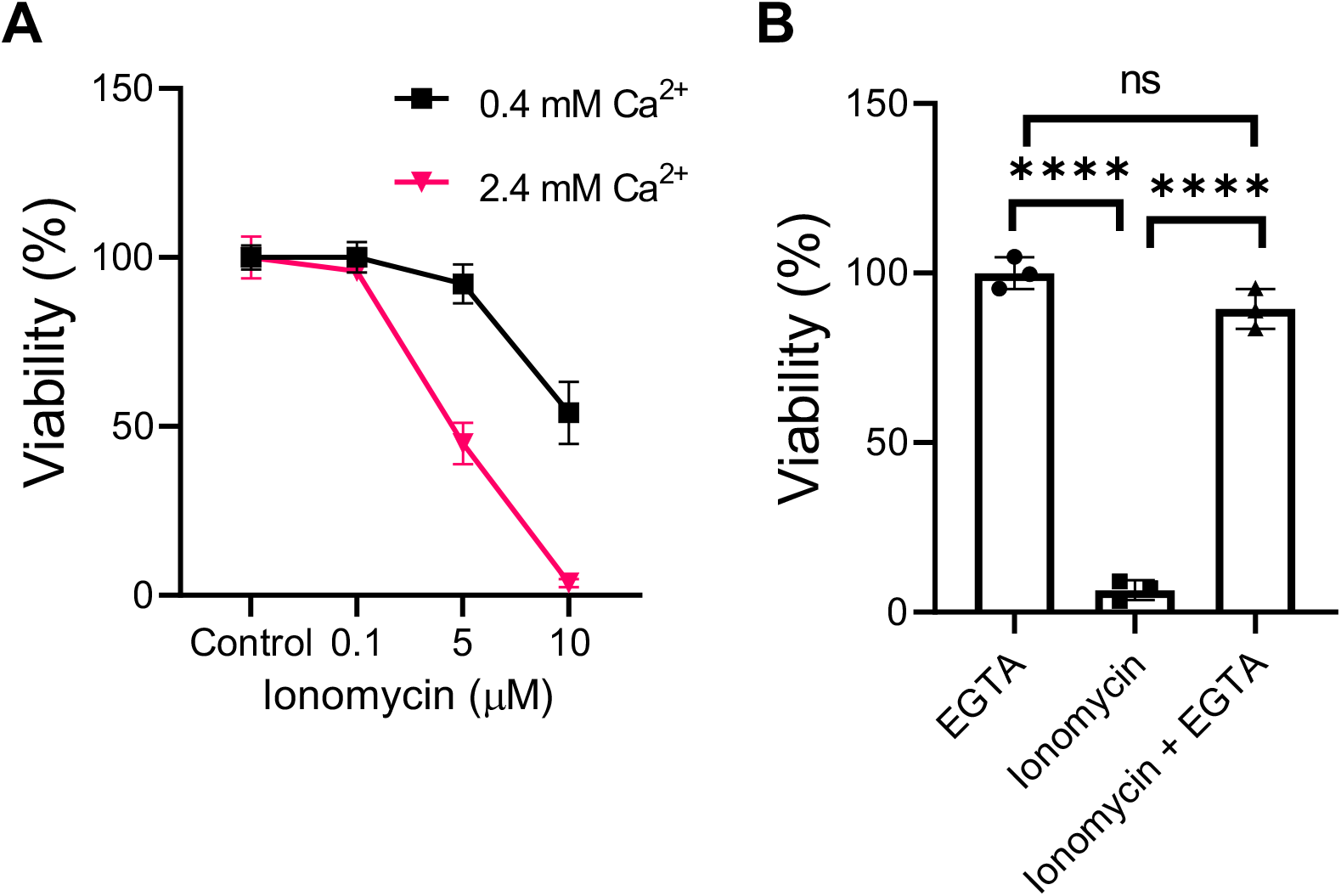
K562 cells are sensitive to Ca^2+^ overload. **(A)** K562 survival after 24 hour incubation with ionomycin in varying Ca^2+^ concentration. **(B)** Ionomycin toxicity is dependent on Ca^2+^. K562 cells incubated with ionomycin for 24 hours with and without Ca^2+^-chelator EGTA (5 mM). Data shown as mean ± SEM. All observations taken from 3 separate biological replicates. ns=p>0.05, ****p<0.0001; significance was determined via one-way ANOVA and Tukey’s multiple comparisons test.

For genome-wide screening, we produced lentivirus from the Brunello library, a validated collection of 76,441 single guide RNAs (sgRNAs) against 19,114 genes across the human genome; each gene is targeted by 4 sgRNAs to include redundancy. Each lentiviral particle encodes for a single gRNA, Cas9 nuclease, and the puromycin resistance gene (Fig. S1A and B). We expanded 144M K562 cells and infected with the Brunello library in triplicate to avoid infection bias. Each infection had ∼48M cells which ensured that each sgRNA was represented by >600 cells in each replicate. This level of coverage was maintained throughout the screen. Importantly, viral transduction was performed at multiplicity of infection (MOI) of 0.3 to ensure that infected cells carry only one viral genome and a single gene knock-out. We manually verified the MOI using a puromycin survival challenge (Fig. S1C). From days 2-10 post infection (dpi), cells underwent puromycin selection to enrich for those carrying the library and to allow time for genomic editing. On day 12, cells were challenged with ionomycin 10 µM or vehicle for 24 hours (Fig. 2A). This led to a 77.6 ± 3.1 % viability loss in treated cells, enriching for cells with protective gene knockouts (Fig. 2B). Surviving cells were allowed to recover and proliferate for 5 days before cell pellet collection and genomic DNA was harvested (Fig. 2B). sgRNA distributions between treated and untreated samples were analyzed via Illumina sequencing at >15M reads per sample (Fig. S1D).

**Figure 2.**
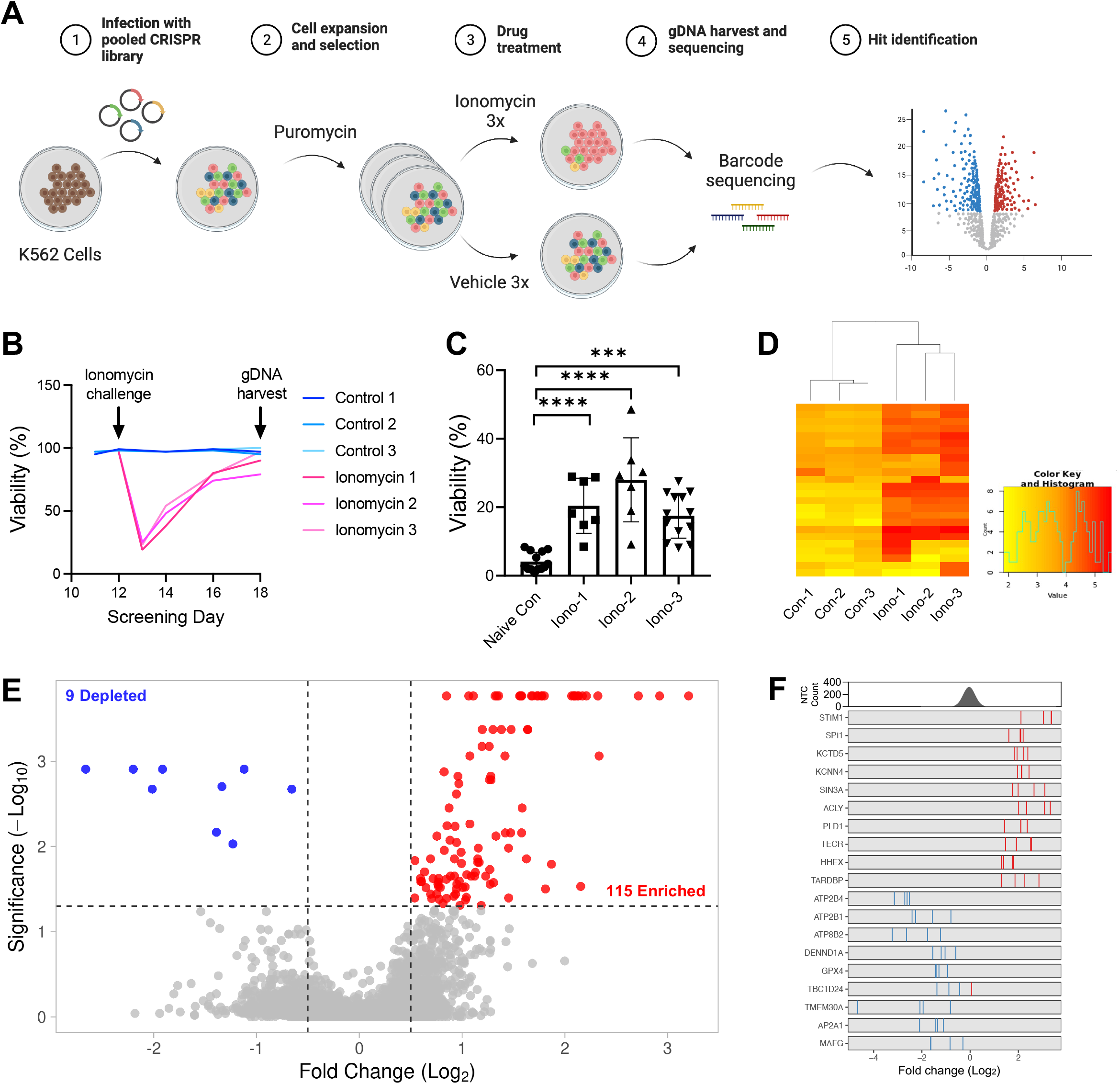
Genome-wide screen identifies cell-autonomous modifiers of Ca^2+^-mediated toxicity. **(A)** CRISPR screening pipeline in K562 cells. CRISPR library infection and ionomycin challenge were done in triplicate. **(B)** Cell viability tracking in screening replicates. Ionomycin 10 µM challenge on day 12 led to massive cell death. Cells were allowed to recover viability past 75% prior to harvest. **(C)** Cell viability following 24-hour ionomycin (7.5 µM) challenge in surviving populations from control cells and ionomycin-challenged cells from *B*. Data points represent technical replicates from three biological replicates. ****p<0.0001, ***p<0.001; significance was determined via one-way ANOVA and Dunnett’s multiple comparisons test. **(D)** Unbiased cluster analysis based on read count variance across samples. **(E)** Volcano plot showing the log2 fold change and false discovery-adjusted P values for all genes following ionomycin challenge. Significantly depleted (blue) and enriched (red) genes defined by a false discovery rate (FDR) <0.05 and fold change >0.5. Non-significant genes shown in grey. **(F)** Fold change distribution of individual sgRNAs for top 10 enriched (blue) and 9 depleted (blue) genes. Frequency distribution of all 1,000 Non-targeting control (NTC) sgRNAs shown above as comparison.

To ascertain that the post-screen cell population was indeed ionomycin-resistant, we treated surviving cells with an additional ionomycin challenge. The surviving cells from the Ca^2+^-challenged arm of the screen were significantly resistant to additional ionomycin compared to control cells (Fig. 2C). Similarly, unbiased cluster analysis on read-count variance across samples demonstrated a clustering of treatment and control conditions (Fig. 2D), suggesting that the challenge was potent enough to diverge variance across both conditions. Lastly, sgRNA dropout was 0.5% and gene drop-out was 0%, indicating that library representation remained stable throughout the screen and no genes were untested (Datafile S1 and S2).

Library reads were analyzed with the validated Model-based Analysis of Genome-wide CRISPR/Cas9 Knockout (MAGeCK) pipeline (see methods). This revealed 115 significantly enriched and 9 depleted genes with a false-discovery cutoff of <0.05 (Fig 2E). Mapping individual sgRNA distributions for top genes against non-targeting controls revealed marked enrichment and depletion (Fig. 2F, Datafile S1 and 2). Reassuringly, top depleted genes included ATP2B1 and ATP2B4, two ATP-dependent calcium pumps known for maintaining Ca^2+^ homeostasis; these serve as internal controls and further support that our screening strategy created sufficient Ca^2+^ overload. Notable among positively enriched genes were STIM1 and ORAI1, two vital players of store operated Ca^2+^ entry (SOCE). Unexpectedly, the screen identified several metabolic enzymes and members of very long chain biosynthesis including ACLY, TECR, HACD2, ELOVL6, and PEDS1. Equally surprising, genetic knockout of various members of the SWItch/Sucrose Non-Fermentable (SWI/SNF) chromatin remodeling machinery—including SMARCD2, SMARCD1, SMARCC1, and CREBP—was protective against Ca^2+^ overload (Fig. 2F, Datafile S1).

Next, we probed multiple gene set databases (KEGG, Gene Ontology, REACTOME, and CORUM) to identify signaling pathways or gene families that were significantly co-enriched or co-depleted in our screen. Notably, fatty acid synthesis and elongation were significantly enriched (Fig. 3A). Similarly, multiple databases identified chromatin remodeling as a significant mediator of Ca^2+^ toxicity (i.e. inhibition provided protection from Ca^2+^), and in particular, the SWI/SNF pathway, histone H3-K27 and K4 methylation, and the Polycomb repressive complex. Lastly, RNA processing was also identified with small nuclear ribonucleoprotein (snRNP) assembly identified by Gene Ontology and Reactome databases independent of each other. In line with those results, spliceosome assembly and mRNA splicing were repeatedly identified by pathway analysis and pointed to the C complex spliceosome and pre-rRNA complex (Fig. 3B). To our knowledge, none of these pathways are explicitly known to modulate sensitivity to Ca^2+^.

**Figure 3.**
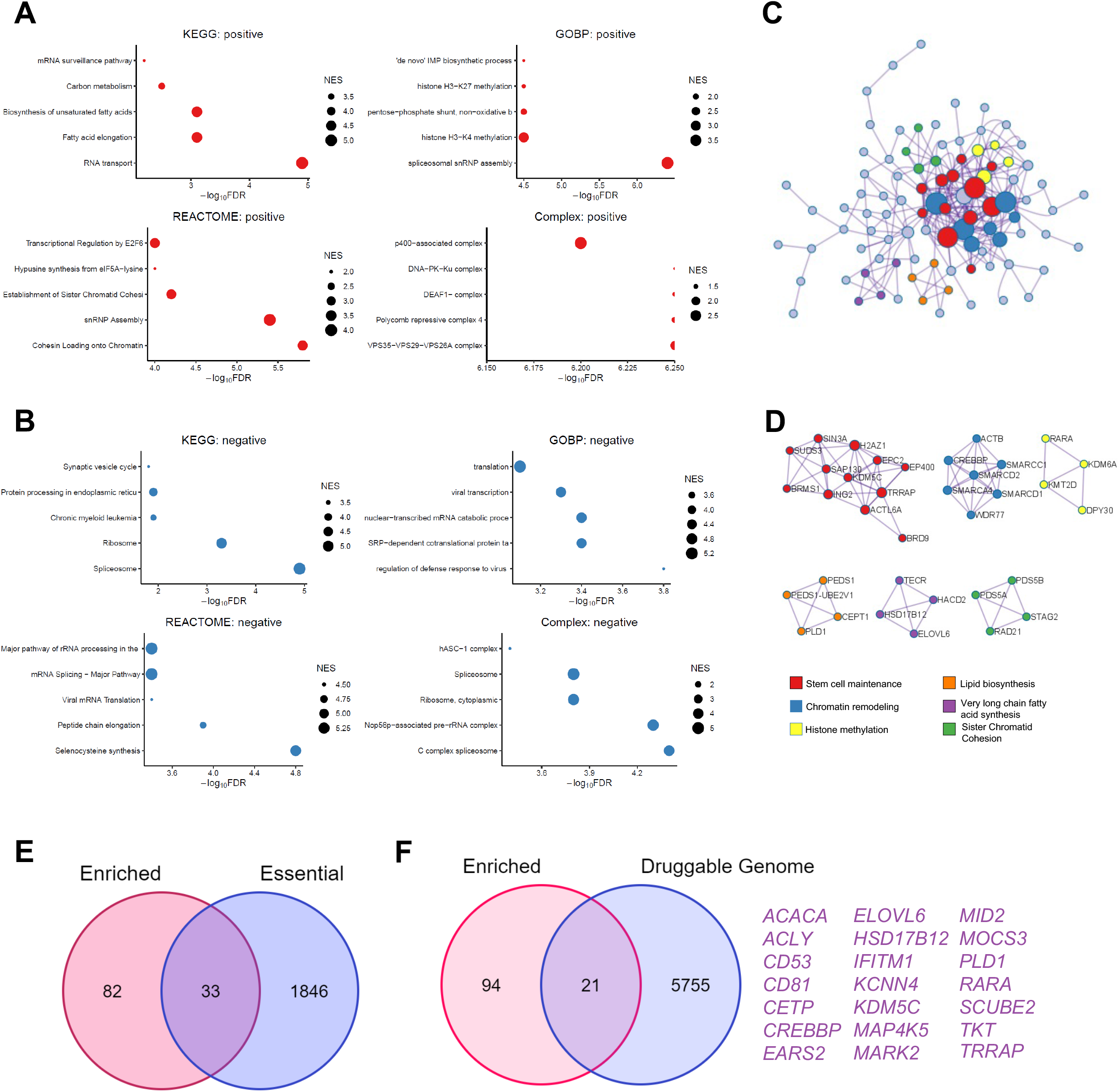
Chromatin remodeling and lipid biosynthesis among enriched pathways. **(A)** Gene set pathway analysis across KEGG, GO, Reactome, and CORUM datasets for enriched and **(B)** depleted genes. NES=Normalized Enrichment Score, FDR=False Discovery Rate-adjusted p value. **(C)** Protein-protein interaction analysis across STRING, BioGrid, OmniPath, InWeb_IM datasets. Color coded gene clusters as in *D*. **(D)** Significantly related subgroups identified and color coded from analysis in C. **(E)** Venn diagram comparing hits enriched in our screen (magenta) versus essential genes (blue) identified in Wang et al. 2015. **(F)** Venn diagram comparing hits enriched in our screen (magenta) versus genes considered part of the druggable genome. All 21 common genes shown in purple.

We then performed protein-interaction analysis to visualize gene clusters that were enriched in our positively enriched hits. This revealed marked interactions among our hits, with hits belonging to stem cell maintenance and chromatin remodeling as central nodes (Fig. 3C). Several sub-cluster families were identified and support the previous pathway analysis with several members of fatty acid biosynthesis, chromatin remodeling, and histone methylation heavily represented (Fig 3D). Our choice of K562 as the screening cell line enabled comparison against previous screens that also utilized this cell line. Comparing our hits against a validated list of essential genes *(37)* (determined in part from K562 cells) reveals that a majority of the genes (82 out of 115) are non-essential (Fig. 3E). Additionally, 21 of our hits are part of the druggable genome *(38)*. Thus, a sizeable portion of our hits represent viable yet unexplored targets for pharmacotherapeutic intervention.

Lastly, we used chemical blockade of the STIM1-ORAI1 axis (both hits in our screen), to determine if their inhibition would protect cells from Ca^2+^ overload. 30-minute pretreatment with ORAI1 inhibitor YM-58483 led to complete rescue of ionomycin-induced cell death (Fig. 4A). Importantly, Ca^2+^-imaging in HEK293T cells showed that YM-58483 does not prevent Ca^2+^ influx due to ionomycin (Fig. 4B).

**Figure 4.**
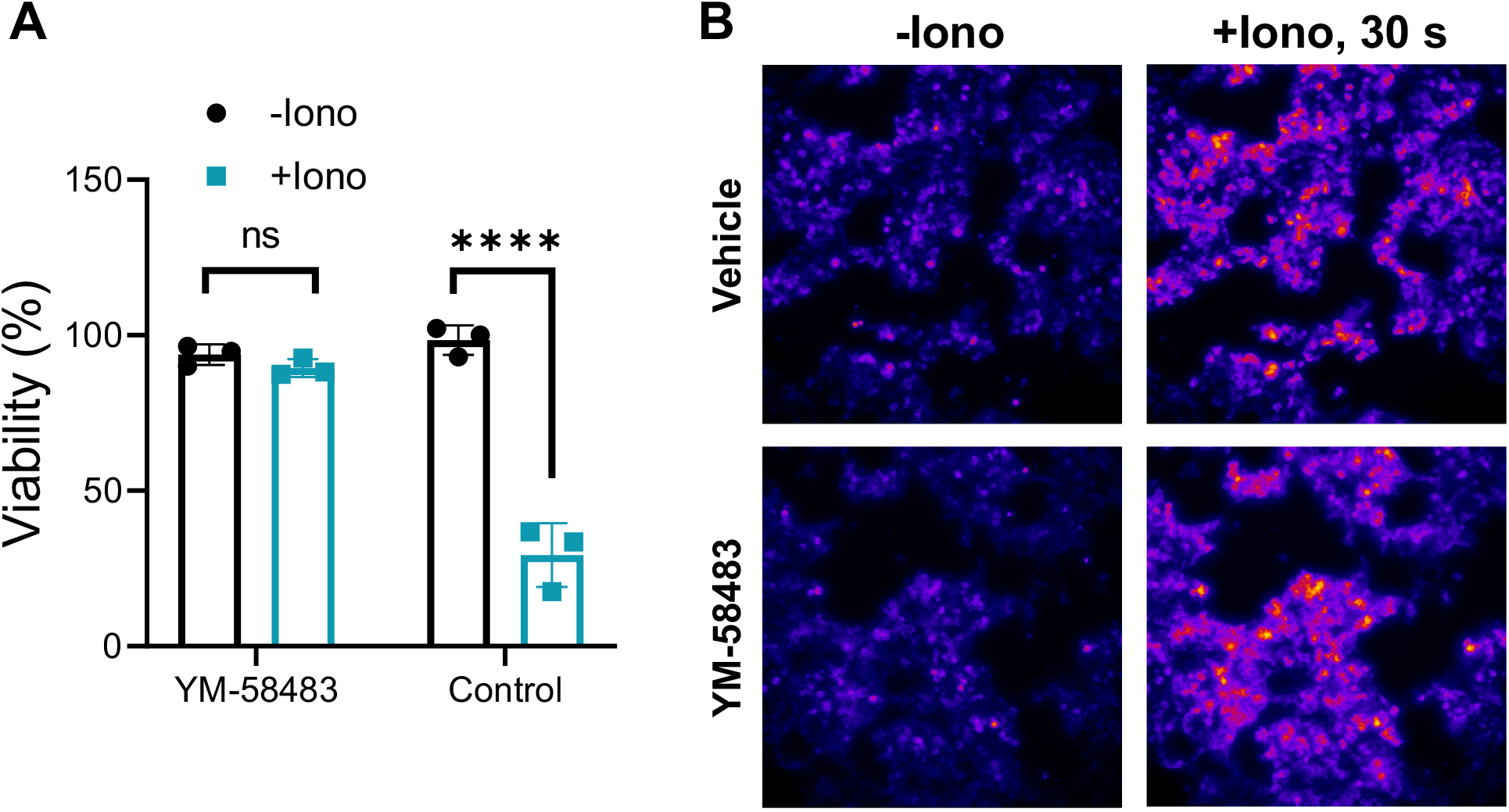
ORAI1 blockade prevents ionomycin-mediated cell death. **(A)** Viability of K562 cells after ionomycin challenge (24 hour, 7.5 µM). Cells were pretreated for 30 minutes with calcium release-activated channel blocker YM-58483 (10 µM). (B) Calcium-imaging in HEK293T cells expressing jRCaMP1a biosensor treated with ionomycin 7.5 µM and ± YM-58483 (10 µM) 30 minute pretreatment. ns=p>0.05, ****p<0.0001; significance was determined via one-way ANOVA and Tukey’s multiple comparisons test.

## DISCUSSION

In this work, we performed a genome-wide assessment for genes that mediate Ca^2+^ toxicity in human cells. We identified K562 cells as a suitable line for this purpose due to their sensitivity to calcium ionophore ionomycin. Using a validated library, we probed the genome and identified 115 protective and 9 deleterious gene knockouts after Ca^2+^ overload. Notably, we identified pathways previously unknown to be involved in Ca^2+^-induced cell death including very long chain fatty acid synthesis, chromatin remodeling, and the SWItch/Sucrose Non-Fermentable (SWI/SNF) complex.

Among enriched hits, we were pleased to identify known modulators of Ca^2+^ entry including STIM1 and ORAI1—both members of SOCE, a pathway previously known to regulate Ca^2+^ in non-excitable cells. Indeed SOCE is a known mediator of apoptosis and neural death after hypoxia *(39–41)*. This validates the ability of our screen to identify bona fide modulators. Our screen identified several other pathways previously unknown to regulate Ca^2+^ toxicity. Interestingly, very long chain fatty acids have been previously known to mediate necroptotic death *(42, 43)*. These studies identified membrane disruption and permeabilization as potential mechanisms but the link of this literature to Ca^2+^ remains unexplored. The SWI/SNF machinery is responsible for chromatin accessibility remodeling, thereby changing genome-wide expression patterns. SWI/SNF activity is known to be pro-survival and its inhibition is being explored for anti-tumor efficacy *(44, 45)*. Thus, it is surprising that its inhibition here led to cell survival; this could hint at context dependent functions and warrant the identification of the chromatin accessibility and expression patterns in SWI/SNF deficient cells—which were protected from Ca^2+^. Encouragingly, for this new role in cancer, a host of small molecule SWI/SNF modulators are emerging *(44, 46)*. These will facilitate exploration of SWI/SNF in contexts where Ca^2+^ is relevant. Lastly, our results identified several genes involved in mRNA handling, including TARDBP, best known for its role in Amyotrophic Lateral Sclerosis (ALS). TARDBP was recently implicated in induction of the mitochondrial permeability transition pore and cell death in neural tissue *(47)*. Similarly, cytosolic Ca^2+^ concentration has been observed to modulate TARDBP localization *(48)*. These results should be further explored given their suggestion that mobilization of TARDBP by Ca^2+^ overload may facilitate TARDP-mediated induction of mitochondrial permeability. Lastly, interactions or interdependence among these pathways is unknown, and an intriguing future direction is to test potential synergy of blocking multiple pathways under Ca^2+^ stress. These findings may serve as a tool for gaining insight into the hierarchy of Ca^2+^-responsive pathways and cell death.

By comparing against the human essential *(37)* and druggable genomes *(38)*, we determined that 82 of the identified protective knockouts were non-essential and 21 are targetable with small molecules. Additionally, we provide proof-of-concept evidence that small molecule blockade of our hits can prevent ionomycin-mediated toxicity. Importantly, ORAI1 inhibition did not appear to impair ionomycin’s ability to import Ca^2+^ into the cell. These data are encouraging, and we are hopeful that small molecule inhibitors targeted against the other non-essential hits could provide similar protection from Ca^2+^ overload in one of the various diseases which feature this mechanism of cell death.

## MATERIALS AND METHODS

### Cell Culture

K562 cells (ATCC CCL-243) and HEK293T/17 cells (ATCC CRL-11268) were maintained in DMEM (L-glutamine, Sodium Pyruvate, Non-essential amino acids; Gibco) supplemented with 10% FBS (Gibco) and Pen/Strep (Gibco). Cells were maintained between 10%-95% confluence (HEK293T) or 100k-1M cells/mL (K562). All cells were maintained in a humidified incubator at 37C with 5% CO_2_. To determine viability, cells were lysed in CellTiter-Glo 2.0 (Promega) reagent and read with a Synergy MX microplate reader. For ORAI1 inhibition, cells were pretreated with YM-58483 (MedChemExpress) or vehicle for 2 hours prior to ionomycin challenge.

### CRISPR screening

The human Brunello CRISPR knockout pooled library was a gift from David Root and John Doench (Addgene #73178) *(49)*. Lentiviral particles were generated in HEK293T as previously described *(31)*. Briefly, HEK293T/17 cells under passage #6 were cultured in 10 cm dishes at 400k cells/mL. These cells were transfected using TransIT-Lenti (Mirus Bio) using a ratio of 5:3.75:1.25 of Brunello:psPAX2:pMD2.G for a total of 10 μg per dish. After 48 hours, supernatant was collected, clarified, and concentrated 10-fold using Lenti-X concentrator (Takara). Lentivirus aliquots were stored at -80 °C until functional titration and use.

Infection into 144M K562 cells was done in triplicate. Cells were spun down for 6 min at 0.2 rcf, then resuspended to 96 mL; 38.4 μL of 10 mg/ml polybrene added (4 μg/mL final), and 960 μL of Brunello virus added (20 μL per well). Mixture was split into 4 12-well plates (2 mL per well) and spun at 1000 rcf for 2 hours at 30C for spinfection. The cells were resuspended and split into 5 t-150 flasks with 50ml at a density of 576k/mL. Viability was tracked daily using CellTiter-Glo 2.0 (Promega) reagent. 2 days after infection, puromycin selection began using 4 μg/mL puromycin. MOI was verified by determining remaining ratio of viable cells at day 4 of puromycin treatment (day at which uninfected control reached complete death). Cells were expanded to ensure >500 cells/sgRNA coverage. Ionomycin 10 μM (or ethanol vehicle) treatment began on day 12 in media containing 2.4 mM Ca^2+^. Treatment media was changed to fresh media after 24 hours. Cells were maintained until pelleting and flash freezing on day 18 post infection. Pellets were ensured to contain >40M cells to maintain sgRNA coverage.

Genomic DNA was extracted using isopropanol precipitation as previously described *(50)*. Cells were digested in a 15 ml conical tube, 6 ml of NK Lysis Buffer (50 mM Tris, 50 mM EDTA, 1% SDS, pH 8) and 30 μl of 20 mg/ml Proteinase K (QIAGEN 19131) were added to the tissue/cell sample and incubated at 55°C overnight. 30 μl of 10 mg/ml RNase A (QIAGEN 19101, diluted in NK Lysis Buffer to 10 mg/ml and then stored at 4°C) was added to the lysed sample, which was then inverted 25 times and incubated at 37°C for 30 min. Genomic DNA was precipitated with 7.5M ammonium acetate and 6 mL 100% isopropyl alcohol. DNA was air dried, resuspended in 0.5 mL TE buffer and stored at -20C.

All genomic DNA from each sample was used for sgRNA amplification with oligos designed for the Brunello library *(49)* using the NEBNext PCR protocol: 2 min at 98C, then 24 cycles: 10 s at 90C, 15 s 69C, 30 s 65C, then 5 min 65C. Each 50 μL reaction contained 2.5 μg of template DNA. Amplified DNA was cleaned up using Monarch DNA clean-up kit (NEB). Amplified DNA was sequenced using single lane NovaSeq: ∼800M reads at 100bp read length.

## Statistics and Bioinformatics

Statistical testing was performed with GraphPad Prism v8.2.0 as described in each figure. NovaSeq sequencing reads were trimmed and compressed using Galaxy Tools. Sequencing data was analyzed with MAGeCK-RRA v0.5.9.4 and visualized with MAGeCKFlute v2.2.0 using default settings *(51, 52)*. Screen quality control was analyzed and visualized with PinAPL-Py v2.9.2 *(53)*.

## Supporting information

Supplemental Datafile

## Acknowledgments

We are thankful to David Mohr and the Johns Hopkins GRCF High Throughput Sequencing Core for expert advice on sequencing analysis. We are thankful to Albert Liu and Rong Li for providing the Brunello Library plasmids.

## Author contributions

Conceptualization: OERG, EDL, MEA

Methodology: OERG

Investigation: OERG Visualization: OERG, KLS

Funding acquisition: OERG, MEA

Project administration: OERG, EDL,

MEA Supervision: OERG, EDL, MEA

Writing – original draft: OERG

Writing – review & editing: OERG, EDL, MEA

