## Supplemental Figures for "Genome-wide CRISPR screen reveals genetic modifiers of Ca^2+^-mediated cell death"

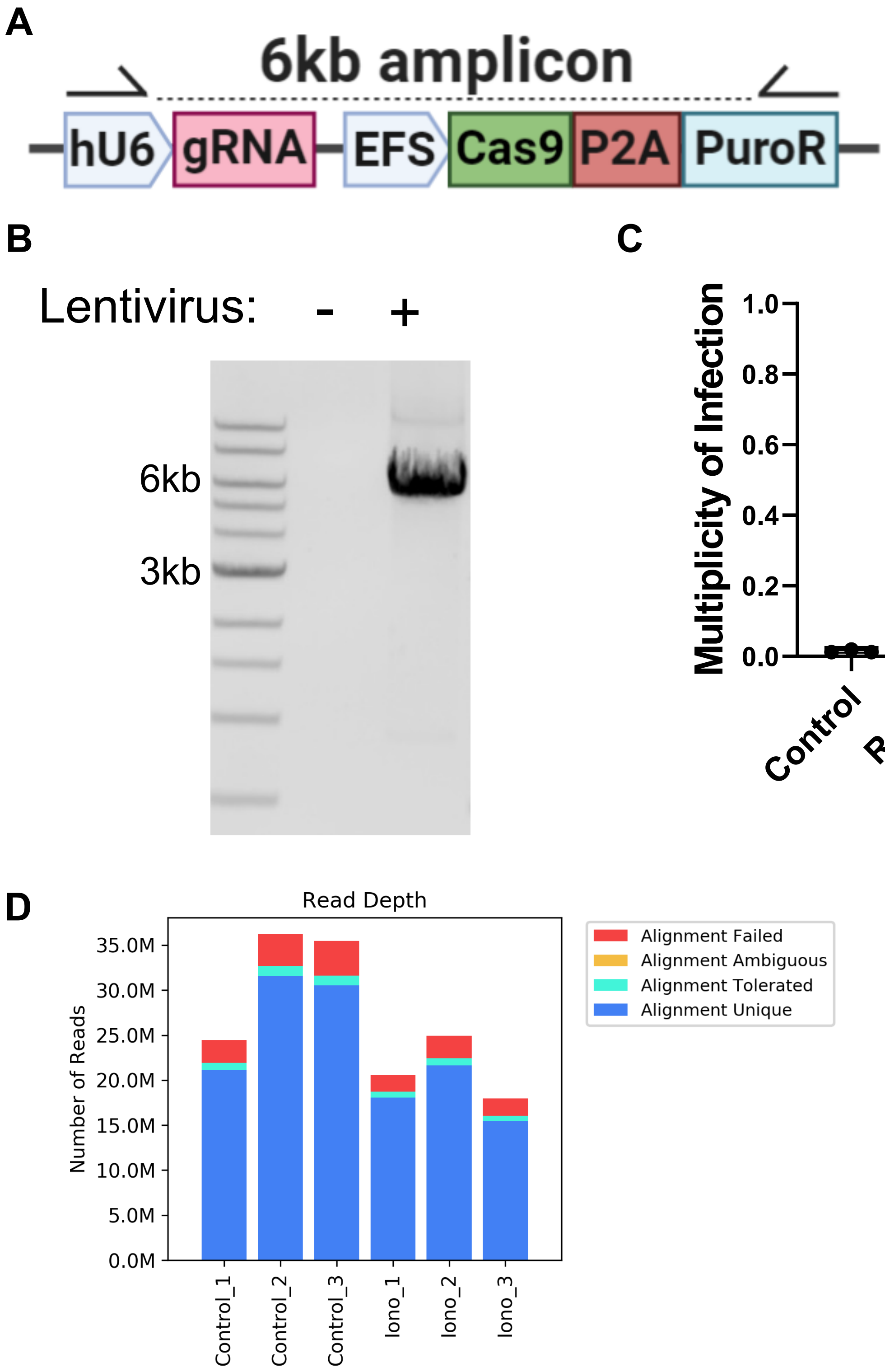


**Figure S1. CRISPR screening quality control. (A)** Schematic of All-In-One lentiviral construct for delivery of Cas9 and sgRNA in Brunello library. **(B)** Integration of CRISPR cassette was verified via PCR from genomic DNA isolated from infected K562 cells. **(C)** Empirical Multiplicity of infection determined via puromycin survival assay in 3 separate replicates used for screening. Data shown as mean ± SEM; mean shown above each bar. **(D)** Sequencing depth and alignment quality in each individual control or ionomycin-challenged (Iono) sample.
